# Effects of reference databases’ refinements on the validity of molecular definitions of 15,371 candidate human-specific regulatory sequences

**DOI:** 10.1101/026435

**Authors:** Gennadi V. Glinsky

## Abstract

Thousands of candidate human-specific regulatory sequences (HSRS) have been identified, supporting the idea that unique to human phenotypes result from human-specific changes to genomic regulatory networks (GRNs). The sequence quality of reference genome databases is essential for the accurate definition of regulatory DNA segments as candidate HSRS. It is unclear how database improvements would affect the validity of the HSRS’ definition. Sequence conservation analysis of 15,371 candidate HSRS was carried out using the most recent releases of reference genomes’ databases of humans and nonhuman primates (NHP) defining the conservation threshold as the minimum ratio of bases that must remap of 1.00. This analysis identifies 3,793 regulatory DNA segments that lack evidence of human-specific mutations and represent regulatory sequences highly conserved in humans, Bonobo, and Chimpanzee. Present analysis revealed a major database refinement’s effect on the validity of HSRS’ definition and suggests that human-specific phenotypes may evolve as a results of integration into human-specific GRNs of both conserved in NHP and human-specific genomic regulatory elements.

## Introduction

Extensive search for human-specific genomic regulatory sequences (HSRS) revealed thousands candidate HSRS, a vast majority of which is residing within non-protein coding genomic regions (McLean et al., 2011; Shulha et al., 2012; Konopka et al., 2012; Capra et al., 2013; Marnetto et al., 2014; Glinsky, 2015). Candidate HSRS comprise multiple distinct families of genomic regulatory elements, which were defined using a multitude of structural features, different statistical algorithms, as well as a broad spectrum of experimental, analytical, computational, and bioinformatics strategies. The current catalogue of candidate HSRS includes conserved in humans novel regulatory DNA sequences designated human accelerated regions, HARs (Capra et al., 2013); fixed human-specific regulatory regions, FHSRR (Marnetto et al., 2014); human-specific transcription factor-binding sites, HSTFBS (Glinsky, 2015), regions of human-specific loss of conserved regulatory DNA termed hCONDEL (McLean et al., 2011); human-specific epigenetic regulatory marks consisting of H3K4me3 histone methylation signatures at transcription start sites in prefrontal neurons (Shulha et al., 2012); and human-specific transcriptional genetic networks in the frontal lobe (Konopka et al., 2012). Most recently, Gittelman et al. (2015) reported identification of 524 DNase I hypersensitive sites (DHSs) that are conserved in nonhuman primates but accelerated in the human lineage (haDHS) and may have contributed to human-specific phenotypes. They estimated that 70% of substitutions in haDHSs are attributable to positive selection consistent with the hypothesis that these DNA segments have been subjects to human-specific adaptive evolution resulting in creation of human-specific regulatory sequences.

Definition of HARs, which is one of the most actively investigated HSRS families, is based on calculations as a baseline the evolutionary expected rate of base substitutions derived from the experimentally determined level of conservation between multiple species at the given locus. The statistical significance of differences between the observed substitution rates on a lineage of interest in relation to the evolutionary expected baseline rate of substitutions can be estimated. This method is considered particularly effective for identifying highly conserved sequences within noncoding genomic regions that have experienced a marked increase of substitution rates on a particular lineage. It has been successfully applied to humans (Pollard et al. 2006; Prabhakar et al. 2006; Bird et al. 2007), where the rapidly-evolving sequences that are highly conserved across mammals and have acquired many sequence changes in humans since divergence from chimpanzees were designated as human accelerated regions (HARs). Experimental analyses of HARs bioactivity revealed that some HARs function as non-coding RNA genes expressed during the neocortex development (Pollard et al., 2006) and human-specific developmental enhancers (Prabhakar et al. 2008). Consistent with the hypothesis that HARs function in human cells as regulatory sequences, most recent computational analyses and transgenic mouse experiments demonstrated that many HARs represent developmental enhancers (Capra et al., 2013).

The high sequence quality of the reference genome databases is essential for accurate definition of regulatory DNA segments as candidate HSRS. To address the problem of a database improvements’ effect on the validity of the HSRS definition, sequence conservation analysis of 15,371 candidate HSRS was carried out using the most recent releases of reference genomes’ databases of humans and nonhuman primates (NHP). In this analysis, the sequence conservation threshold was defined as the minimum ratio of bases that must remap of 1.00. Present analysis demonstrates that a total of 3,793 regulatory DNA segments (Supplemental Data Set 1), which were previously defined as candidate HSRS, should be classified as the highly conserved in humans and nonhuman primates regulatory sequences, thus reducing the number of candidate HSRS by 24.7%.

## Results and Discussion

### Sequence conservation analysis of human accelerated DNase I hypersensitive sites

The identified haDHSs represent relatively short DNA segments of the median size 290 bp. (range from 150- 1010 bp.; average size of 323 bp.), which are predominantly located within intronic and intergenic sequences (Gittelman et al., 2015). To test whether reported 524 haDHSs represent human-specific DNA sequences, the conservation analysis was carried-out using the LiftOver algorithm and Multiz Alignments of 20 mammals (17 primates) of the UCSC Genome Browser on Human Dec. 2013 (GRCh38/hg38) Assembly (http://genome.ucsc.edu/cgi-bin/hgTracks?db=hg38&position=chr1%3A90820922-90821071&hgsid=441235989_eelAivpkubSY2AxzLhSXKL5ut7TN).

The most recent releases of the corresponding reference genome databases were utilized to ensure the use of the most precise, accurate, and reproducible genomic DNA sequences available to date. The results of these analyses are reported in the Table 1. Several thresholds of the LiftOver algorithm MinMatch function (minimum ratio of bases that must remap) were utilized to assess the sequences conservation and identify candidate human-specific regulatory sequences as previously described (Glinsky, 2015). In this analysis, the candidate human-specific regulatory sequences were defined based on conversion failures to both Chimpanzee’s and Bonobo’s genomes and supported by direct visual evidence of human-specific sequence alignment differences of the Multiz Alignments of 20 mammals (17 primates). It appears that only small fractions (0.2%-13.9%) of reported 524 haDHSs can be defined as candidate human-specific regulatory sequences even using different sequence conservation thresholds (Table 1). Based on this analysis, the vast majority (86.1% to 99.8%) of 524 haDHSs could be classified as the candidate regulatory sequences that appear conserved in humans and nonhuman primates. Therefore, a major reference genome database refinements’ effect was observed on definition of haDHS family of candidate HSRS.

**Table 1.** Distribution of primate-specific and human-specific regulatory sequences among 524 haDHSs reported by Gittelman et al. (2015).

| **Genomes/LiftOver setting** | **MinMatch 0.95** | **MinMatch 0.99** | **MinMatch 1.00** |
| --- | --- | --- | --- |
| **Human genome (hg19)** | 524 | 524 | 524 |
| **Human genome (hg38)** | 524 | 524 | 524 |
| **Mouse genome conversion (mm10)** | 298 | 148 | 66 |
| **Percent conserved in rodents' genome** | 56.9 | 28.2 | 12.6 |
| **Chimpanzee genome conversion** | 520 | 493 | 439 |
| **Percent conserved in Chimpanzee** | 99.2 | 94.1 | 83.8 |
| **Chimpanzee conversion failures\*** | 4 | 31 | 85 |
| **Bonobo genome conversion** | 517 | 492 | 425 |
| **Percent conserved in Bonobo** | 98.7 | 93.9 | 81.1 |
| **Bonobo conversion failures** | 7 | 30 | 99 |
| **Human-specific sequences\*\*** | **1** | **21** | **73** |
| **Percent conserved in non-human primates** | **99.8** | **96.0** | **86.1** |
| **Percent of human-specific sequences** | **0.2** | **4.0** | **13.9** |

Legends: LiftOver algorithm MinMatch, Minimum ratio of bases that must remap; haDHS, human accelerated DNase I hypersensitive sites;
\*Chimpanzee genome PanTro4 conversion;
\*\*Human-specific regulatory sequences were defined based on conversion failures to both Chimpanzee and Bonobo genomes

Interestingly, the Multiz Alignments of 20 mammals (17 primates) revealed that 71% of candidate human-specific haDHSs defined at 0.99 MinMatch threshold (Table 1) contain small human-specific inserts of 2-15 bp., suggesting a common mutation mechanism (Supplemental Table S1). A majority (78%) of candidate human-specific haDHSs are located within the intronic (47.9%) and intergenic (30.1%) sequences (Supplemental Table S2). However, 15 of 73 (20.5%) candidate human-specific haDHSs sequences appear to intersect exons, 11 of which include intron/exon junctions (Supplemental Tables S1& S2). Intriguingly, this analysis identified the 18 bp. human-specific deletion within the exon 9 of the *PAX8* gene, which appears to affect the structure of the *PAX8-AS1* RNA as well (Supplemental Table S1).

### Sequence conservation analysis of human accelerated regions

Strikingly similar results were observed when the sequence conservation analysis of 2,745 HARs was performed (Table 2). It appears that only small fractions (1.2%-9.3%) of reported HARs can be defined as candidate human-specific regulatory sequences using different sequence conservation thresholds (Table 2). Based on this analysis, the vast majority (90.7% to 98.8%) of 2,745 HARs could be classified as the candidate regulatory sequences that appear conserved in humans and nonhuman primates (Table 2). Therefore, there is a major reference genome database refinements’ effect on the accuracy of definition of the HARs family of candidate HSRS.

**Table 2.** Distribution of primate-specific and human-specific regulatory sequences among 2,741 HARs reported by Capra et al. (2013).

| **Genomes/LiftOver setting** | **MinMatch 0.95** | **MinMatch 0.99** | **MinMatch 1.00** |
| --- | --- | --- | --- |
| **Human genome (hg19)** | 2,745 | 2,745 | 2,745 |
| **Human genome (hg38)** | 2,741 | 2,740 | 2,739 |
| **Mouse genome conversion (mm10)** | 2,364 | 1,642 | 1,004 |
| **Percent conserved in rodents' genome** | 86.2 | 59.9 | 36.7 |
| **Chimpanzee genome conversion** | 2,698 | 2,608 | 2,404 |
| **Percent conserved in Chimpanzee** | 98.4 | 95.2 | 87.8 |
| **Chimpanzee conversion failures\*** | 43 | 133 | 337 |
| **Bonobo genome conversion** | 2,687 | 2,590 | 2,341 |
| **Percent conserved in Bonobo** | 98.0 | 94.5 | 85.5 |
| **Bonobo conversion failures** | 54 | 151 | 400 |
| **Human-specific sequences\*\*** | **33** | **107** | **255** |
| **Percent conserved in non-human primates** | **98.8** | **96.1** | **90.7** |
| **Percent of human-specific sequences** | **1.2** | **3.9** | **9.3** |

Legends: LiftOver algorithm MinMatch, Minimum ratio of bases that must remap; HARs, human accelerated regions;
\*Chimpanzee genome PanTro4 conversion;
\*\*Human-specific regulatory sequences were defined based on conversion failures to both Chimpanzee and Bonobo genomes

### Sequence conservation analysis of other classes of candidate HSRS

In contrast to haDHS and HARs, other classes of candidate HSRS were defined based on the failure of alignments of human regulatory DNA segments to the reference genome databases of other species (Marnetto et al., 2014; Glinsky, 2015). It appears that a majority (82.1%-88.4%) of reported DNase I hypersensitive sites-derived fixed human specific regulatory regions (DHS_FHSRR) can be defined as candidate human-specific regulatory sequences using different sequence conservation thresholds (Table 3). Based on this analysis, the relatively minor fraction (11.6% to 17.9%) of 2,118 DHS_FHSRR may be classified as the candidate regulatory sequences that appear conserved in humans and nonhuman primates (Table 3).

**Table 3.** Distribution of primate-specific and human-specific regulatory sequences among 2,118 DHS fixed human specific regulatory regions reported by Marnetto et al. (2014).

| **Genomes/LiftOver setting** | **MinMatch 0.95** | **MinMatch 0.99** | **MinMatch 1.00** |
| --- | --- | --- | --- |
| **Human genome (hg19)** | 2,118 | 2,118 | 2,118 |
| **Human genome (hg38)** | 2,116 | 2,115 | 2,114 |
| **Mouse genome conversion (mm10)** | 18 | 18 | 4 |
| **Percent conserved in rodents' genome** | 0.9 | 0.9 | 0.2 |
| **Chimpanzee genome conversion** | 5 | 5 | 5 |
| **Percent conserved in Chimpanzee** | 0.2 | 0.2 | 0.2 |
| **Chimpanzee conversion failures\*** | 2,111 | 2,111 | 2,111 |
| **Bonobo genome conversion** | 375 | 331 | 242 |
| **Percent conserved in Bonobo** | 17.7 | 15.7 | 11.4 |
| **Bonobo conversion failures** | 1,741 | 1,785 | 1,874 |
| **Human-specific sequences\*\*** | **1,737** | **1,781** | **1,869** |
| **Percent conserved in non-human primates** | **17.9** | **15.8** | **11.6** |
| **Percent of human-specific sequences** | **82.1** | **84.2** | **88.4** |

Legends: LiftOver algorithm MinMatch, Minimum ratio of bases that must remap; DHS, DNase I hypersensitive sites;
\*Chimpanzee genome PanTro4 conversion;
\*\*Human-specific regulatory sequences were defined based on conversion failures to both Chimpanzee and Bonobo genomes

Similarly, a majority (79.0%-86.5%) of reported HSTFBS can be defined as candidate human-specific regulatory sequences using different sequence conservation thresholds (Table 4). The relatively minor fraction (13.5% to 21.0%) of 3,803 HSTFBS may be classified as the candidate regulatory sequences that appear conserved in humans and nonhuman primates (Table 4). A majority (70.2%-79.7%) of reported hESC_FHSRR can be defined as candidate human-specific regulatory sequences using different sequence conservation thresholds (Table 5). The relatively small fraction (20.3% to 29.8%) of 1,932 hESC_FHSRR could be classified as the candidate regulatory sequences that appear conserved in humans and nonhuman primates (Table 6). A majority (84.3%-89.7%) of reported other_FHSRR can be defined as candidate human-specific regulatory sequences using different sequence conservation thresholds (Table 5). The relatively minor fraction (10.3% to 15.7%) of 4,249 other_FHSRR could be classified as the candidate regulatory sequences that appear conserved in humans and nonhuman primates (Table 6). Based on this analysis, it appears that there is a relatively minor reference genome database refinements’ effect on the accuracy of molecular definition of HSTFBS and FHSRR families of candidate HSRS.

**Table 4.** Distribution of primate-specific and human-specific regulatory sequences among 3,803 human specific transcription factor-binding sites reported by Glinsky (2015).

| **Genomes/LiftOver setting** | **MinMatch 0.95** | **MinMatch 0.99** | **MinMatch 1.00** |
| --- | --- | --- | --- |
| **Human genome (hg19)** | 3,803 | 3,803 | 3,803 |
| **Human genome (hg38)** | 3,719 | 3,714 | 3,714 |
| **Mouse genome conversion (mm10)** | 31 | 13 | 12 |
| **Percent conserved in rodents' genome** | 0.8 | 0.4 | 0.3 |
| **Chimpanzee genome conversion** | 70 | 60 | 56 |
| **Percent conserved in Chimpanzee** | 1.9 | 1.6 | 1.5 |
| **Chimpanzee conversion failures\*** | 3,649 | 3,659 | 3,663 |
| **Bonobo genome conversion** | 768 | 529 | 495 |
| **Percent conserved in Bonobo** | 20.7 | 14.2 | 13.3 |
| **Bonobo conversion failures** | 2,951 | 3,190 | 3,224 |
| **Human-specific sequences\*\*** | **2,937** | **3,173** | **3,211** |
| **Percent conserved in non-human primates** | **21.0** | **14.6** | **13.5** |
| **Percent of human-specific sequences** | **79.0** | **85.4** | **86.5** |

Legends: LiftOver algorithm MinMatch, Minimum ratio of bases that must remap;
\*Chimpanzee genome PanTro4 conversion;
\*\*Human-specific regulatory sequences were defined based on conversion failures to both Chimpanzee and Bonobo genomes

**Table 5.** Distribution of primate-specific and human-specific regulatory sequences among 1,932 hESC fixed human specific regulatory regions reported by Marnetto et al. (2014).

| **Genomes/LiftOver setting** | **MinMatch 0.95** | **MinMatch 0.99** | **MinMatch 1.00** |
| --- | --- | --- | --- |
| **Human genome (hg19)** | 1,932 | 1,932 | 1,932 |
| **Human genome (hg38)** | 1,932 | 1,930 | 1,928 |
| **Mouse genome conversion (mm10)** | 0 | 0 | 0 |
| **Percent conserved in rodents' genome** | 0.0 | 0.0 | 0.0 |
| **Chimpanzee genome conversion** | 1 | 1 | 0 |
| **Percent conserved in Chimpanzee** | 0.1 | 0.1 | 0.0 |
| **Chimpanzee conversion failures\*** | 1,931 | 1,931 | 1,932 |
| **Bonobo genome conversion** | 575 | 529 | 396 |
| **Percent conserved in Bonobo** | 29.8 | 27.4 | 20.5 |
| **Bonobo conversion failures** | 1,357 | 1,403 | 1,536 |
| **Human-specific sequences\*\*** | **1,357** | **1,403** | **1,536** |
| **Percent conserved in non-human primates** | **29.8** | **27.3** | **20.3** |
| **Percent of human-specific sequences** | **70.2** | **72.7** | **79.7** |

Legends: LiftOver algorithm MinMatch, Minimum ratio of bases that must remap; hESC, human embryonic stem cells;
\*Chimpanzee genome PanTro4 conversion;
\*\*Human-specific regulatory sequences were defined based on conversion failures to both Chimpanzee and Bonobo genomes

**Table 6.**
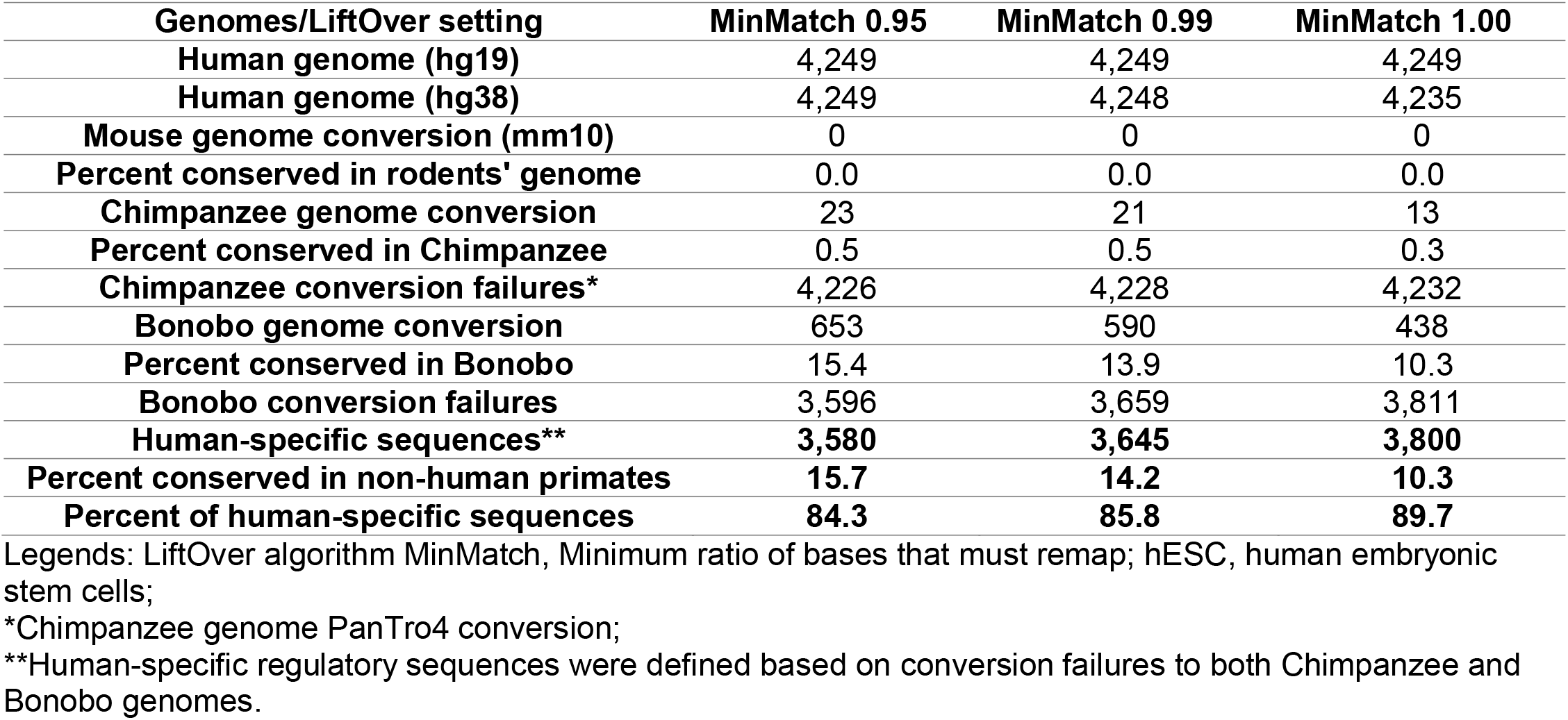
Distribution of primate-specific and human-specific regulatory sequences among 4,249 fixed human specific regulatory regions reported by Marnetto et al. (2014).

### Identification of highly conserved in nonhuman primates regulatory DNA sequences among candidate HSRS

To identify regulatory DNA segments that are highly conserved in nonhuman primates, the most stringent definition of the sequence conservation threshold was used by setting the minimum sequence alignments’ match requirement as the ratio of bases that must remap of 1.00. Significantly, a given regulatory DNA segment was defined as highly conserved only when both direct and reciprocal conversions between humans’ and nonhuman primates’ genomes were observed using the MinMatch threshold of 1.00. Strikingly, the majority of both haDHSs (404 of 524; 77.1%) and HARs (2,262 of 2,739; 82.6%) were defined as the highly conserved in humans and NHP regulatory sequences lacking evidence of human-specific mutations. In contrast, only relatively small fractions of other classes of candidate HSRS were identified as highly conserved in nonhuman primates regulatory sequences, scored at 7.3% for HSTFBS; 8.3% for FHSRR; 9.4% for DHS_FHSRR; and 15.9% for hESC_FHSRR (Table 7). Based on the results of the present analysis (Table 7), the majority of HARs and haDHSs cannot be classified as the candidate HSRS; they should be defined as the regulatory sequences that are highly conserved in humans and nonhuman primates and lack evidence of human-specific mutations. A total of 3,793 regulatory DNA segments (Supplemental Data Set 1), which were previously defined as candidate HSRS, should be classified as the highly conserved in humans and nonhuman primates regulatory sequences, thus reducing the number of candidate HSRS by 24.7%.

**Table 7.**
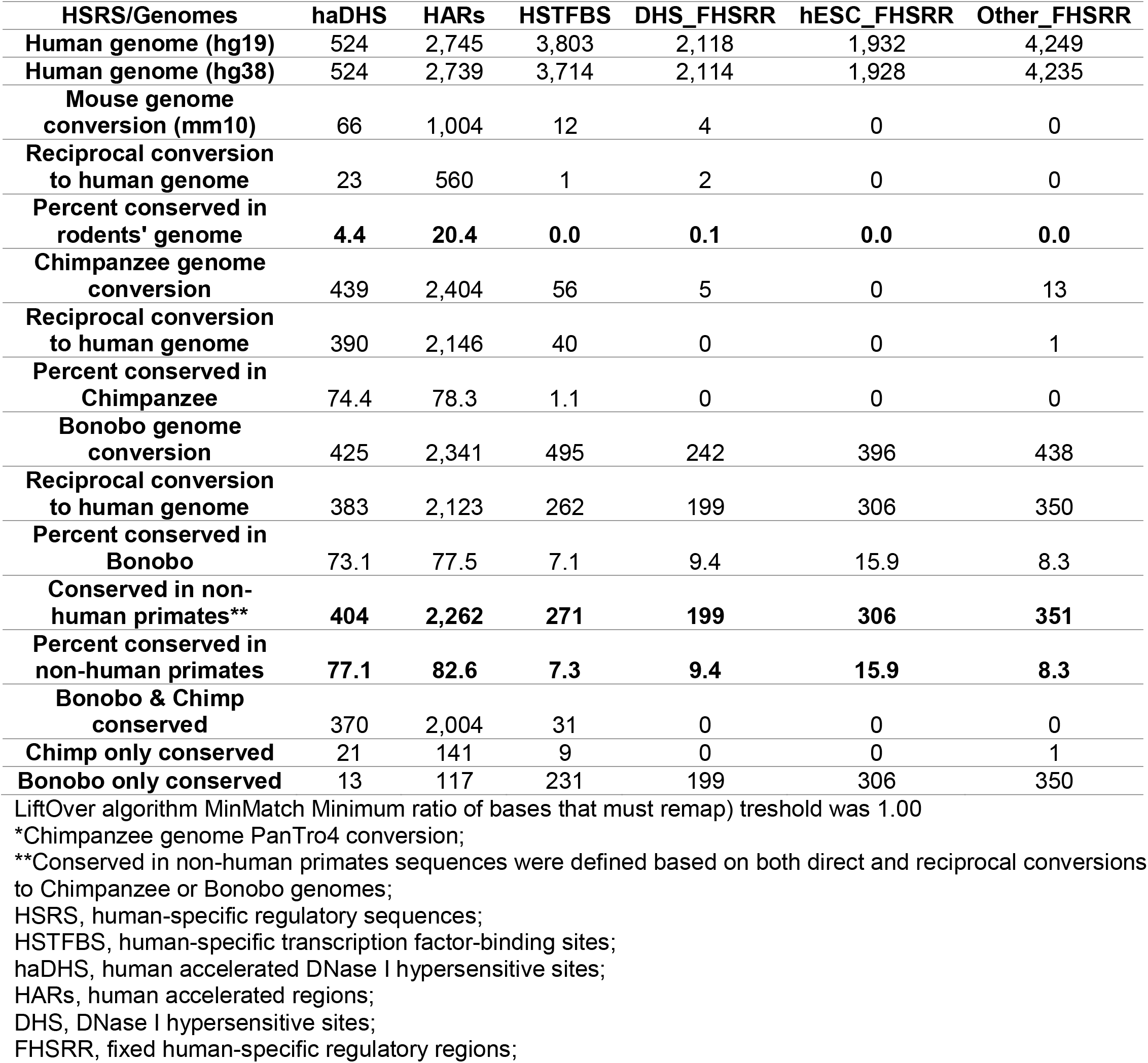
Distribution of highly conserved in non-human primates regulatory sequences among 15,371 candidate human-specific regulatory sequence.

| **HSRS/Genomes** | **haDHS** | **HARs** | **HSTFBS** | **DHS_FHSRR** | **hESC_FHSRR** | **Other_FHSRR** |
| --- | --- | --- | --- | --- | --- | --- |
| **Human genome (hg19)** | 524 | 2,745 | 3,803 | 2,118 | 1,932 | 4,249 |
| **Human genome (hg38)** | 524 | 2,739 | 3,714 | 2,114 | 1,928 | 4,235 |
| **Mouse genome conversion (mm10)** | 66 | 1,004 | 12 | 4 | 0 | 0 |
| **Reciprocal conversion to human genome** | 23 | 560 | 1 | 2 | 0 | 0 |
| **Percent conserved in rodents' genome** | **4.4** | **20.4** | **0.0** | **0.1** | **0.0** | **0.0** |
| **Chimpanzee genome conversion** | 439 | 2,404 | 56 | 5 | 0 | 13 |
| **Reciprocal conversion to human genome** | 390 | 2,146 | 40 | 0 | 0 | 1 |
| **Percent conserved in Chimpanzee** | 74.4 | 78.3 | 1.1 | 0 | 0 | 0 |
| **Bonobo genome conversion** | 425 | 2,341 | 495 | 242 | 396 | 438 |
| **Reciprocal conversion to human genome** | 383 | 2,123 | 262 | 199 | 306 | 350 |
| **Percent conserved in Bonobo** | 73.1 | 77.5 | 7.1 | 9.4 | 15.9 | 8.3 |
| **Conserved in non-human primates\*\*** | **404** | **2,262** | **271** | **199** | **306** | **351** |
| **Percent conserved in non-human primates** | **77.1** | **82.6** | **7.3** | **9.4** | **15.9** | **8.3** |
| **Bonobo & Chimp conserved** | 370 | 2,004 | 31 | 0 | 0 | 0 |
| **Chimp only conserved** | 21 | 141 | 9 | 0 | 0 | 1 |
| **Bonobo only conserved** | 13 | 117 | 231 | 199 | 306 | 350 |

LiftOver algorithm MinMatch Minimum ratio of bases that must remap) threshold was 1.00
\*Chimpanzee genome PanTro4 conversion;
\*\*Conserved in non-human primates sequences were defined based on both direct and reciprocal conversions to Chimpanzee or Bonobo genomes;
HSRS, human-specific regulatory sequences;
HSTFBS, human-specific transcription factor-binding sites;
haDHS, human accelerated DNase I hypersensitive sites;
HARs, human accelerated regions;
DHS, DNase I hypersensitive sites;
FHSRR, fixed human-specific regulatory regions;

### Conclusions

Present analysis revealed a major reference database refinement’s effect on the validity of molecular definitions of candidate HSRS. A total of 3,793 regulatory DNA segments, which were previously defined as candidate HSRS, appear lacking evidence of human-specific mutations and represent regulatory sequences highly conserved in humans, Bonobo, and Chimpanzee. Reported herein sequence conservation analysis reveals that only a small fraction of haDHS and HARs loci can be defined as candidate HSRS. A majority of haDHSs and HARs appear to represent highly conserved in humans and nonhuman primates candidate regulatory sequences, suggesting that human-specific phenotypes may evolve as a result of combinatorial interplay of both conserved in nonhuman primates and human-specific regulatory sequences.

## Methods

### Data source

Solely publicly available datasets and resources were used for this analysis. A total of 15,371 candidate HSRS were analyzed in this study, including 2,745 human accelerated regions (Capra et al., 2013), 524 human accelerated DNase I hypersensitive sites (Gittelman et al., 2015), 3,083 human-specific transcription factor binding sites (Glinsky, 2015), and 8,229 fixed human-specific regulatory regions, FHSRR (Marnetto et al., 2014) divided into 2,118 DHS_FHSRR; 1,932 hESC_FHSRR; and 4,249 FHSRR identified in different human cell lines, excluding hESC (Other_FHSRR).

### Data analysis

To test whether reported 15,371 candidate HSRS represent human-specific regulatory DNA sequences, the conservation analysis was carried-out using the LiftOver algorithm and Multiz Alignments of 20 mammals (17 primates) of the UCSC Genome Browser (Kent et al., 2002) on Human Dec. 2013 Assembly (GRCh38/hg38) (http://genome.ucsc.edu/cgi-bin/hgTracks?db=hg38&position=chr1%3A90820922-90821071&hgsid=441235989_eelAivpkubSY2AxzLhSXKL5ut7TN). The most recent releases of the corresponding reference genome databases were utilized to ensure the use of the most precise, accurate, and reproducible genomic DNA sequences available to date. A candidate HSRS was considered conserved if it could be aligned to either one or both Chimpanzee or Bonobo genomes using defined sequence conservation thresholds of the LiftOver algorithm MinMatch function. LiftOver conversion of the coordinates of human blocks to non-human genomes using chain files of pre-computed whole-genome BLASTZ alignments with a specified MinMatch levels and other search parameters in default setting (http://genome.ucsc.edu/cgi-bin/hgLiftOver). Several thresholds of the LiftOver algorithm MinMatch function (minimum ratio of bases that must remap) were utilized to assess the sequences conservation and identify candidate human-specific (MinMatch of 0.95; 0.99; and 1.00) and conserved in nonhuman primates (MinMatch of 1.00) regulatory sequences as previously described (Glinsky, 2015). The Net alignments provided by the UCSC Genome Browser were utilized to compare the sequences in the human genome (hg38) with the mouse (mm10), Chimpanzee (PanTro4), and Bonobo genomes. A given regulatory DNA segment was defined as the highly conserved regulatory sequence when both direct and reciprocal conversions between humans’ and nonhuman primates’ genomes were observed using the MinMatch sequence alignment threshold of 1.00 (Table 7). A given regulatory DNA segment was defined as the candidate human-specific regulatory sequence when sequence alignments failed to both Chimpanzee and Bonobo genomes using the specified MinMatch sequence alignment thresholds (Tables 1-6).

## Supplemental Information

Supplemental information includes Supplemental Tables S1 and S2; Supplemental Data Set 1; and can be found with this article online.

## Author Contributions

This is a single author contribution. All elements of this work, including the conception of ideas, formulation, and development of concepts, execution of experiments, analysis of data, and writing of the paper, were performed by the author.

## Acknowledgements

This work was made possible by the open public access policies of major grant funding agencies and international genomic databases and the willingness of many investigators worldwide to share their primary research data. I would like to thank my anonymous colleagues for their valuable critical contributions during the peer review process of this work.

